# Cholesterol remodels the endoplasmic reticulum to control myofibroblastic CAF function

**DOI:** 10.64898/2026.02.16.706237

**Authors:** Srimayee Vaidyanathan, Nicole L Yuwono, Ellie TY Mok, Jessica L Chitty, Jesse A Rudd-Schmidt, Riley A Goldsworthy, Rasan M. Sathiqu, Andrew G Cox, Thomas R Cox, Kristin K Brown

## Abstract

Chronic exposure of local fibroblasts to the reactive tumour niche results in the emergence of cancer-associated fibroblasts (CAFs), which can be classed into multiple subtypes with distinct functional and molecular characteristics. Of these subtypes, myofibroblastic CAFs (myCAFs), contribute to tumour progression primarily by depositing and remodelling extracellular matrix (ECM) within the tumour microenvironment (TME). ECM production is known to be a metabolically demanding process, but specific metabolic pathways that are reprogrammed during myCAF activation are poorly understood. Here, we show that cholesterol biosynthesis is elevated in myCAFs relative to normal fibroblasts. We find that cholesterol is specifically enriched in the myCAF endoplasmic reticulum (ER) membrane and is associated with augmented ER mass and enhanced ER secretory function. Importantly, we demonstrate that pharmacological or genetic inhibition of cholesterol biosynthesis plays a critical role in regulating the ECM-secreting and ECM remodelling function of myCAFs. Moreover, we show that inhibition of cholesterol biosynthesis in myCAFs markedly constrains mammary tumour metastasis *in vivo*. These studies suggest that cholesterol biosynthesis represents a metabolic vulnerability that can be exploited to normalise myCAF-mediated ECM dysfunction in the TME.

## Introduction

Dysfunction of the local stromal compartment is a feature inherent to most solid tumours. Long-standing clinical observations have described the transition from hyperplasia to malignancy as being accompanied by the recruitment, expansion and activation of resident fibroblast populations (1). Chronic exposure of these local fibroblasts to the reactive tumour niche results in the emergence of cancer-associated fibroblasts (CAFs), which represent a major cellular component in malignancies such as breast and pancreatic carcinoma (2). CAFs can be classed into several subtypes, each with distinct functional and molecular characteristics (2–4). Of these subtypes, myofibroblastic CAFs (myCAFs) are characterized by phenotypic similarity to myofibroblasts generated during wound-healing and fibrosis. myCAFs demonstrate an augmented capacity for depositing and modifying collagenous extracellular matrix (ECM), thus promoting a fibrosis-like response known as desmoplasia. In breast carcinomas, altered ECM architecture is not just a feature of tumorigenesis but also a driver, with breast density having been recognized for a number of decades as a risk factor for malignant transformation (5,6). Desmoplastic reorganization of ECM by myCAFs actively contributes to tumour progression by modulating several known hallmarks of cancers. For instance, invasion and metastasis are promoted by the radial alignment of collagen fibres at the tumour-stroma interface, as well as the production of matrix-remodelling enzymes, which create permissive tracks in the tumour ECM (7,8). Increased ECM crosslinking and stiffening also enhance tumour progression through the induction of mechanosensitive oncogenic signalling cascades (9,10). Altered stromal mechanics and density can additionally impede the efficacy of a range of cytotoxic therapies (11–13), as well as promote immune cell exclusion (14,15).

Although targeting myCAF-mediated desmoplasia represents an attractive therapeutic strategy, attempts to do so have yielded mixed results. For instance, while targeting ECM crosslinking using lysyl oxidase (LOX) inhibitors has been shown to effectively decrease desmoplasia in a number of cancer types (13,16), complete ablation of myCAFs or myCAF-derived collagen paradoxically increases tumour cell invasion (17,18). This highlights the need for a more detailed understanding of the specific molecular pathways underpinning the ECM-producing and ECM-remodelling functions of myCAFs to identify more nuanced approaches to target the tumour-promoting functions of myCAFs.

The production of desmoplastic ECM by myCAFs represents a significant anabolic demand, with the process of collagen synthesis, secretion and modification containing multiple nodes of potential regulation by metabolites and metabolic pathways. Indeed, several recent studies support the notion that CAF function and aberrant ECM deposition are intrinsically underpinned by rewiring of cellular metabolism (19–21). Given that several processes critical to the translation, modification and secretion of collagens and other ECM-associated proteins are localized within the ER, establishment of the myCAF state poses significant pressure upon the ER machinery. However, little is known regarding the role of ER remodelling and metabolic adaptation to ER stress in myCAFs compared with their normal counterparts.

In this study, we demonstrate that cholesterol biosynthesis is upregulated in a mammary myCAF model, leading to cholesterol enrichment within the ER membrane. Importantly, we find that ER mass and function are increased in myCAFs, and that this phenomenon is dependent on cholesterol biosynthesis, as well as the XBP1-mediated arm of the unfolded protein response (UPR). Finally, we demonstrate that genetic and pharmacological inhibition of cholesterol biosynthesis suppresses ER secretory function, collagen deposition and ECM-remodelling capacity. Importantly, we find that inhibition of cholesterol biosynthesis in myCAFs can inhibit the metastatic potential of breast cancer cells *in vivo*. We thus identify cholesterol-dependent remodelling of the ER as a metabolic vulnerability that can be exploited to normalize myCAF function.

## Results

### CAFs exhibit reprogrammed cholesterol metabolism and enrichment of cholesterol within the ER membrane

CAFs isolated from mammary tumours arising in transgenic MMTV-PyMT mice have been described to most closely resemble the myCAF subtype, exhibiting increased expression of alpha-smooth muscle actin (αSMA), as well as increased collagen I deposition and ECM-remodelling capacity relative to normal mammary fibroblasts (NFs) (22). To more effectively study fibroblast metabolism *in vitro,* fibroblasts were cultured in the physiological media Plasmax (23). RNA-Seq analysis revealed elevated expression of multiple myCAF (herein referred to as CAF) markers, including *Acta2* (αSMA) and *Fn1* (fibronectin), when CAFs were cultured in Plasmax versus conventional medium (DMEM) (**Supplementary Figure 1A**). Consistent with the observed transcriptional changes, αSMA and fibronectin expression was also augmented at the protein level in Plasmax (**Supplementary Figure 1B**). Having established that culture in physiological media yields a more robust CAF signature, all further analyses were carried out with fibroblasts cultured in Plasmax.

To gain insight into metabolic differences existing between NFs and CAFs, cells were labelled with [U-^14^C]glucose and fractionated into the five major classes of macromolecules, namely polar metabolites, DNA, RNA, non-polar metabolites (predominantly lipids) and protein (24) (**Supplementary Figure 1C**). While glucose uptake was comparable between the two cell types (**Supplementary Figure 1D**), a significant increase in incorporation of glucose-derived carbon into lipids, and concomitant decrease in incorporation into RNA, was observed in CAFs relative to NFs (**Figure 1A**). To more specifically examine differences in lipid metabolism between NFs and CAFs, cells were labelled with [1-^14^C]acetate, an alternative carbon source for lipid synthesis. Consistent with results from the [U-^14^C]glucose labelling studies, incorporation of [1-^14^C]acetate into lipids was significantly increased in CAFs indicating increased lipid synthesis in CAFs relative to NFs (**Figure 1B**). Interestingly, RNA-Seq analysis revealed a pronounced enrichment of a lipid metabolism signature in CAFs compared with NFs (**Figure 1C**). The cellular lipid pool comprises various species, including sterols, fatty acids, and fatty acyl-derived species such as glycerolipids, phospholipids and sphingolipids. Interestingly, robust upregulation of genes contributing to cholesterol biosynthesis was observed in CAFs (**Figure 1D**). Elevated expression of the rate-limiting enzyme of cholesterol biosynthesis, 3-hydroxy-3-methylglutaryl-CoA reductase (HMGCR) was confirmed in CAFs at the level of protein expression (**Supplementary Figure 1E**). Consistent with these findings, total cholesterol levels were elevated in CAFs relative to NFs (**Figure 1E**). Analysis of published single cell RNA-Seq (scRNA-Seq) data generated from 26 primary patient-derived breast tumours (25) demonstrated that genes encoding the cholesterol biosynthetic enzymes squalene epoxidase (*SQLE*) and sterol-C5-desaturase (*SC5D*) are enriched within the myCAF-like cluster, indicating that cholesterol biosynthesis is a feature of the mammary myCAF phenotype in patients (**Figure 1F**). Interestingly, expression of *SQLE* in the stromal compartment has previously been associated with poor clinical prognosis (26). Cholesterol is a critical component of cellular membranes, where it modulates rigidity and integrity (27). In contrast to NFs, where filipin staining revealed localisation of free cholesterol in large droplets throughout the cytoplasm, accumulation of free cholesterol was observed in the perinuclear region of CAFs (**Figure 1G**). The localisation and distribution of filipin staining in CAFs was suggestive of cholesterol enrichment within the ER membrane where cholesterol levels are typically low (28). In support of this finding, increased abundance of free cholesterol was detected in ER fractions isolated from CAFs (**Figure 1H** and **Supplementary Figure 1F**). Together, these data suggest that reprogramming of cholesterol biosynthesis is a metabolic feature of the CAF state and is associated with increased cholesterol abundance within the ER membrane.

**Figure 1:**
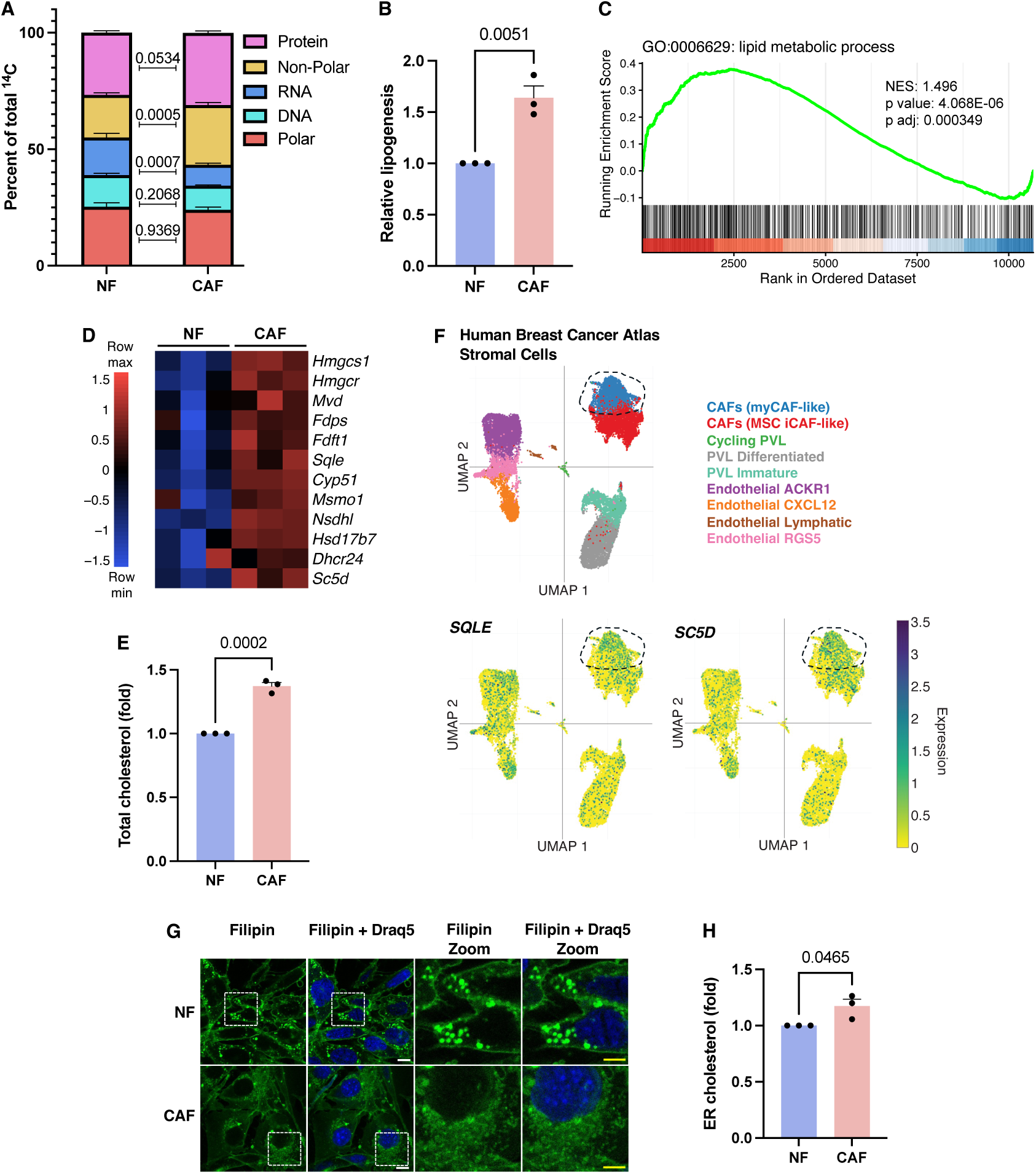
CAFs exhibit reprogrammed cholesterol metabolism and enrichment of cholesterol within the ER membrane. **(A)** Measurement of the relative incorporation of [U-^14^C]glucose into protein, non-polar, RNA, DNA and polar fractions following 24 hours of labelling, n=3. **(B)** Relative incorporation of [1-^14^C]acetate into the lipid fraction of NFs and CAFs, n=3. **(C)** Gene Set Enrichment Analysis (GSEA) plot derived from RNA-Seq analysis of CAFs versus NFs, showing enrichment of a “lipid metabolic process” signature (GO:0006629). **(D)** Heatmap of cholesterol biosynthesis gene expression in NFs and CAFs, as determined by RNA-Seq analysis, n=3. **(E)** Quantification of relative total cholesterol levels in NFs and CAFs, n=3. **(F)** UMAP visualisation of 19,601 patient-derived breast tumour stromal cells analysed by single cell RNA-Seq (scRNA-Seq) from the Human Breast Cancer Atlas (25). Stromal clusters corresponding to immunomodulatory CAFs (iCAFs), myCAFs, perivascular (PVL) and endothelial cell subsets are indicated by colour (top). Feature plots illustrating expression of *SQLE* (bottom left) and *SC5D* (bottom right) in breast tumour stromal cell clusters. Gene expression is represented by log-normalised expression values. The myCAF cluster is delineated by a dashed line. **(G)** Representative filipin staining (green) of free cholesterol in NFs and CAFs. Nuclei are stained with Draq5 (blue). Dashed boxes indicate magnified regions shown to the right. White scale bar represents 10 µm, yellow scale bar represents 5 µm. **(H)** Quantification of relative cholesterol levels in the ER-enriched fraction isolated from NFs and CAFs, n=3.

### CAFs exhibit augmented ER sheet mass and function

Given that ECM proteins are synthesised and processed in the ER, CAFs would be expected to be highly dependent on a functional ER network. Indeed, gene set enrichment analysis (GSEA) revealed upregulation of a gene signature associated with the ER membrane in CAFs relative to NFs (**Figure 2A**). To further examine alterations to the ER network in CAFs, total ER mass was assessed by ER-Tracker staining. Consistent with transcriptomic analysis, a striking increase in total ER mass was observed in CAFs relative to NFs (**Figure 2B**). The ER is a network of interconnected sheets and tubules, with ribosome-studded ER sheets being a site of protein synthesis and secretion, and smooth ER tubules being a site of lipid synthesis. To determine whether ER expansion in CAFs was consistent across both ER sheets and ER tubules, immunofluorescent (IF) staining was performed for the sheet marker cytoskeleton-associated protein 4 (CKAP4) and the tubule marker reticulon-4 (RTN4). Whereas a robust increase in CKAP4-positive ER sheets was observed in CAFs relative to NFs, there was no change in RTN4-positive ER tubules (**Figure 2C**). Interestingly, increased accumulation of the ER exit site (ERES) protein transport and Golgi organization 1 (TANGO1), which facilitates export of large cargo such as procollagens (29), was also observed in CAFs (**Figure 2D)**.

**Figure 2:**
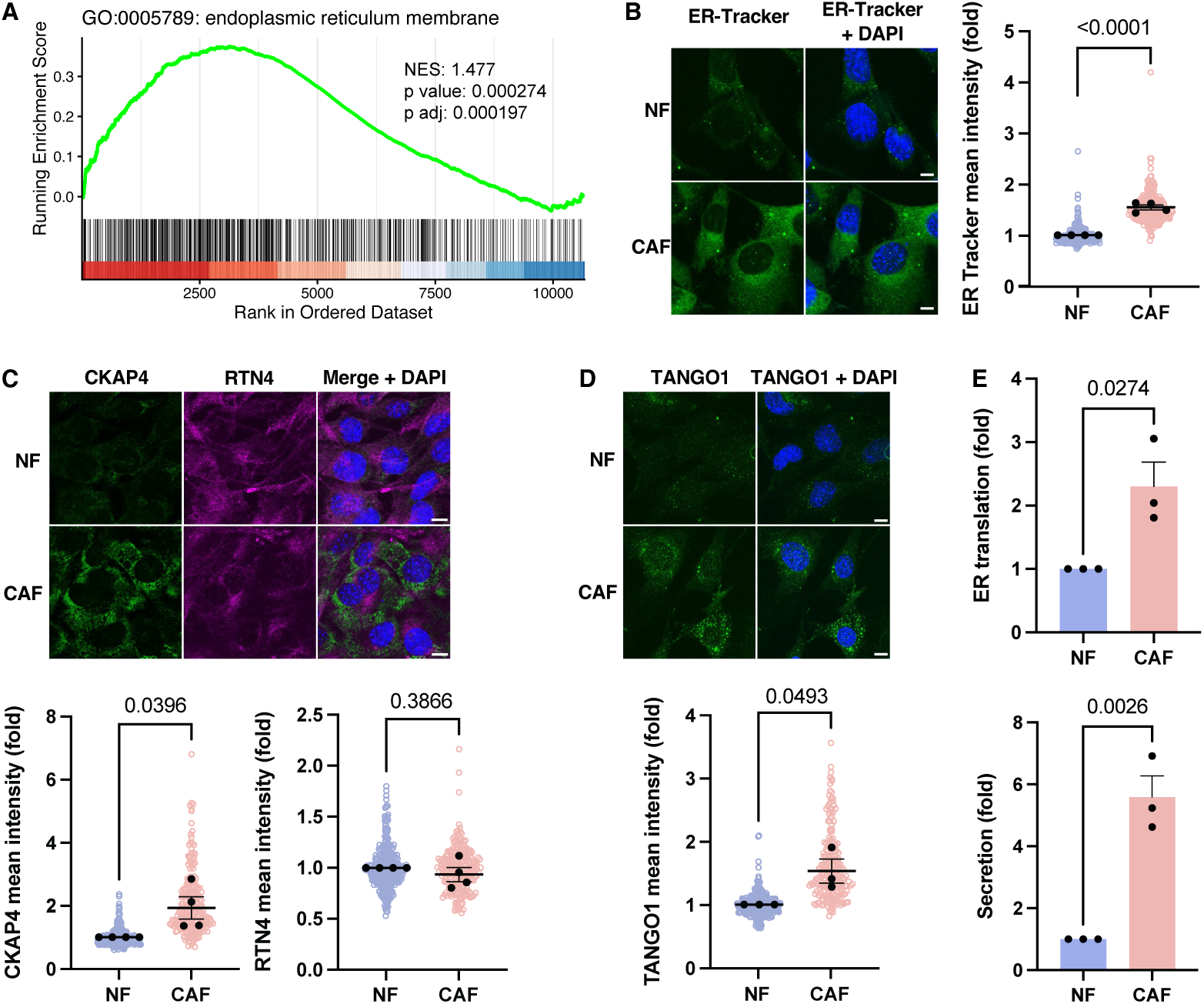
CAFs exhibit augmented ER sheet mass and function. **(A)** GSEA plot derived from RNA-Seq analysis of CAFs versus NFs, showing enrichment of an “endoplasmic reticulum membrane” signature (GO:0005789). **(B)** Representative ER-Tracker staining (green) in NFs and CAFs, and quantification of ER-Tracker mean intensity per cell, n=4 independent experiments (NF=317 cells, CAF=240 cells). Nuclei are stained with DAPI (blue). Scale bars represent 10 µm. **(C)** Representative immunofluorescent staining of ER sheet marker CKAP4 (green) and ER tubule marker RTN4 (magenta) in NFs and CAFs, and quantification of CKAP4 and RTN4 mean intensity per cell, n=4 independent experiments (NF=340 cells, CAF=225 cells). Nuclei are stained with DAPI (blue). Scale bars represent 10 µm. **(D)** Representative immunofluorescent staining of TANGO1 in NFs and CAFs, and quantification of TANGO1 mean intensity per cell, n=3 independent experiments (NF=295 cells, CAF=214 cells). Nuclei are stained with DAPI (blue). Scale bars represent 10 µm. **(E)** Relative ER translation capacity (intracellular EGFP fluorescence intensity) and relative ER secretory capacity (extracellular GLuc activity) in CAFs versus NFs expressing GLuc-T2A-EGFP, n=3.

Given that ER sheets represent a critical compartment at which all secreted and transmembrane proteins are translated, folded, modified and packaged for transport, we postulated that expansion of ER sheet mass in CAFs may facilitate increased synthesis and secretion of ECM proteins and matrix-remodelling factors by these cells. To assess ER sheet function, NFs and CAFs were transduced with a lentiviral construct expressing the naturally secreted Gaussia luciferase (GLuc) protein (30) and EGFP, separated by a T2A self-cleaving peptide (GLuc-T2A-EGFP). As GLuc contains an N-terminal signal peptide, both proteins are translated at the rough ER. Intracellular EGFP fluorescence and extracellular GLuc activity, surrogate markers of ER translation and secretory capacity respectively, were significantly increased in CAFs relative to NFs (**Figure 2E**). Collectively, these findings suggest that ER mass and ER secretory function are increased in CAFs relative to NFs.

### XBP1 regulates ER function in CAFs

Increased ER biogenesis is one of several processes elicited downstream of the unfolded protein response (UPR) to increase the protein-folding capacity of the ER and mitigate ER stress. Given the observed increase in ER mass and secretory function exhibited by CAFs, we next investigated the activation state of the UPR in NFs and CAFs. Increased expression of target genes associated with the inositol-requiring enzyme 1 (IRE1)/X-box-binding protein 1 (XBP1)-mediated arm of the UPR was observed in CAFs relative to NFs (**Figure 3A**). Processing of *Xbp1* mRNA by active IRE1 is required to generate the *Xbp1s* splice isoform, which encodes the active XBP1 transcription factor (31,32). Both total *Xbp1* and *Xbp1s* were elevated in CAFs relative to NFs (**Figure 3B)**. To investigate a mechanistic role for XBP1 in regulating CAF ER function, genetic depletion of *Xbp1* was performed (**Supplementary Figure 2A**). *Xbp1* knockdown induced a significant decrease in both ER translation and secretory capacity in CAFs but had no impact on NFs (**Figure 3C**), suggesting that CAF ER function is specifically sensitive to loss of XBP1. Interestingly, the reduction in ER translation and secretory capacity was accompanied by a dramatic reduction in ER-Tracker staining (**Figure 3D**). Depletion of *Xbp1* also resulted in increased staining with thioflavin T (ThT), a dye that exhibits enhanced fluorescence upon binding to misfolded protein aggregates and therefore serves as a marker of ER dysfunction (**Figure 3E**). Having determined that CAF ER function is at least partially dependent on XBP1, we next investigated whether *Xbp1* depletion impacts the ECM-secreting and ECM-remodelling function of CAFs. While *Xbp1* depletion did not significantly impact collagen I production (**Supplementary Figure 2B**), a significant inhibition of CAF ECM-remodelling capacity was observed in the context of *Xbp1* knockdown (**Figure 3F**). Together these data suggest that XBP1 contributes to the regulation of ER function in CAFs.

**Figure 3:**
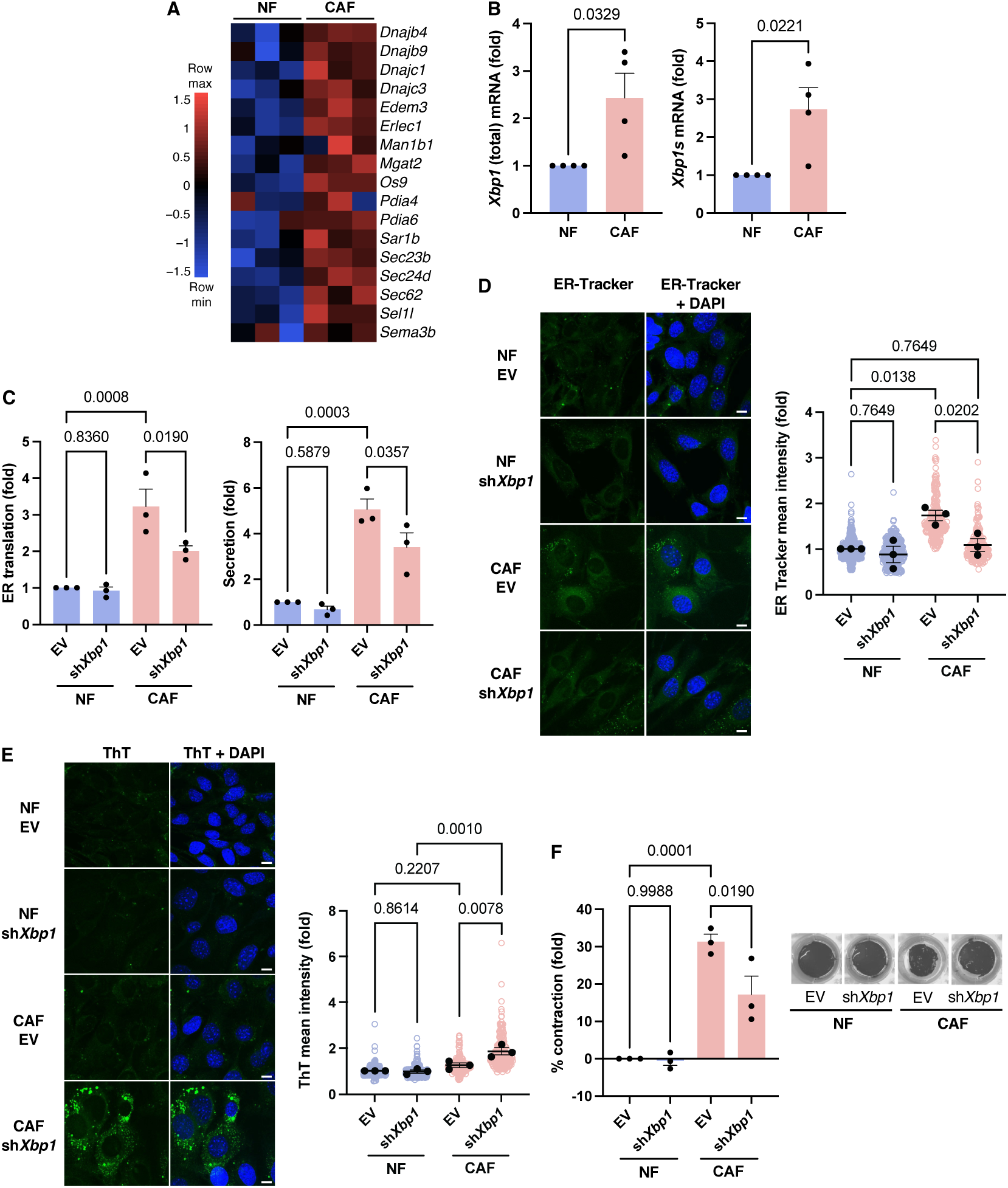
XBP1 regulates ER function in CAFs. **(A)** Heatmap of XBP1 target gene expression in NFs and CAFs, as determined by RNA-seq analysis, n=3. **(B)** Expression of total *Xbp1* mRNA (left) and spliced *Xbp1* (*Xbp1s*) mRNA (right) in NFs versus CAFs, as determined via qPCR, n=4. **(C)** Relative ER translation capacity (intracellular EGFP fluorescence intensity) and relative ER secretory capacity (extracellular GLuc activity) in GLuc-T2A-EGFP-expressing NFs and CAFs transduced with empty vector (EV) or sh*Xbp1* constructs, n=3. **(D)** Representative ER-Tracker staining (green) in NFs and CAFs transduced with EV or sh*Xbp1* constructs, and quantification of ER-Tracker mean intensity per cell, n=3 independent experiments (NF EV=554 cells, NF sh*Xbp1*=375 cells, CAF EV=273 cells, CAF sh*Xbp1*=251 cells). Nuclei are stained with DAPI (blue). Scale bars represent 10 µm. **(E)** Representative thioflavin T (ThT) staining in NFs and CAFs transduced with EV or sh*Xbp1* constructs, and quantification of ThT intensity per cell, n=3 independent experiments (NF EV=478 cells, NF sh*Xbp1*=395 cells, CAF EV=256 cells, CAF sh*Xbp1*=231 cells). Nuclei are stained with DAPI (blue). Scale bars represent 10 µm. **(F)** ECM remodelling capacity of NFs and CAFs transduced with EV or sh*Xbp1* constructs, as determined by a collagen contraction assay over 24 hours. Data are presented as % contraction relative to NF EV, n=3.

### CAF ER function is dependent on cholesterol biosynthesis

Given the observed increase in ER cholesterol in CAFs (**Figure 1H**), we next sought to determine whether cholesterol biosynthesis contributes to altered ER function in CAFs. Previous studies have suggested that cholesterol is particularly abundant at ERES and is necessary for the export of secretory cargo from the ER (33,34). Interestingly, pharmacological inhibition of HMGCR with simvastatin led to a reduction in ER-Tracker staining and CKAP4-positive ER sheet mass in CAFs (**Figure 4A,B**). Moreover, in a subset of simvastatin-treated CAFs, CKAP4-stained ER appeared to be highly fragmented suggesting that acute inhibition of cholesterol biosynthesis disrupts ER sheet architecture (**Figure 4B**). Consistent with perturbed ER sheet mass, pharmacological inhibition of HMGCR disrupted the ER secretory capacity of CAFs without impacting translation capacity (**Figure 4C**). Moreover, simvastatin treatment suppressed TANGO1 accumulation (**Figure 4D**) and increased ThT staining (**Figure 4E**) in CAFs, indicating a loss of competent ERESs and elevated protein misfolding respectively. Results observed following acute pharmacological inhibition of HMGCR were recapitulated following genetic depletion of *Hmgcr* (**Supplementary Figure 3A-E**). Overall, these data reveal that reprogrammed cholesterol biosynthesis contributes to augmented ER sheet mass and ERES function in CAFs, thereby supporting protein secretion.

**Figure 4:**
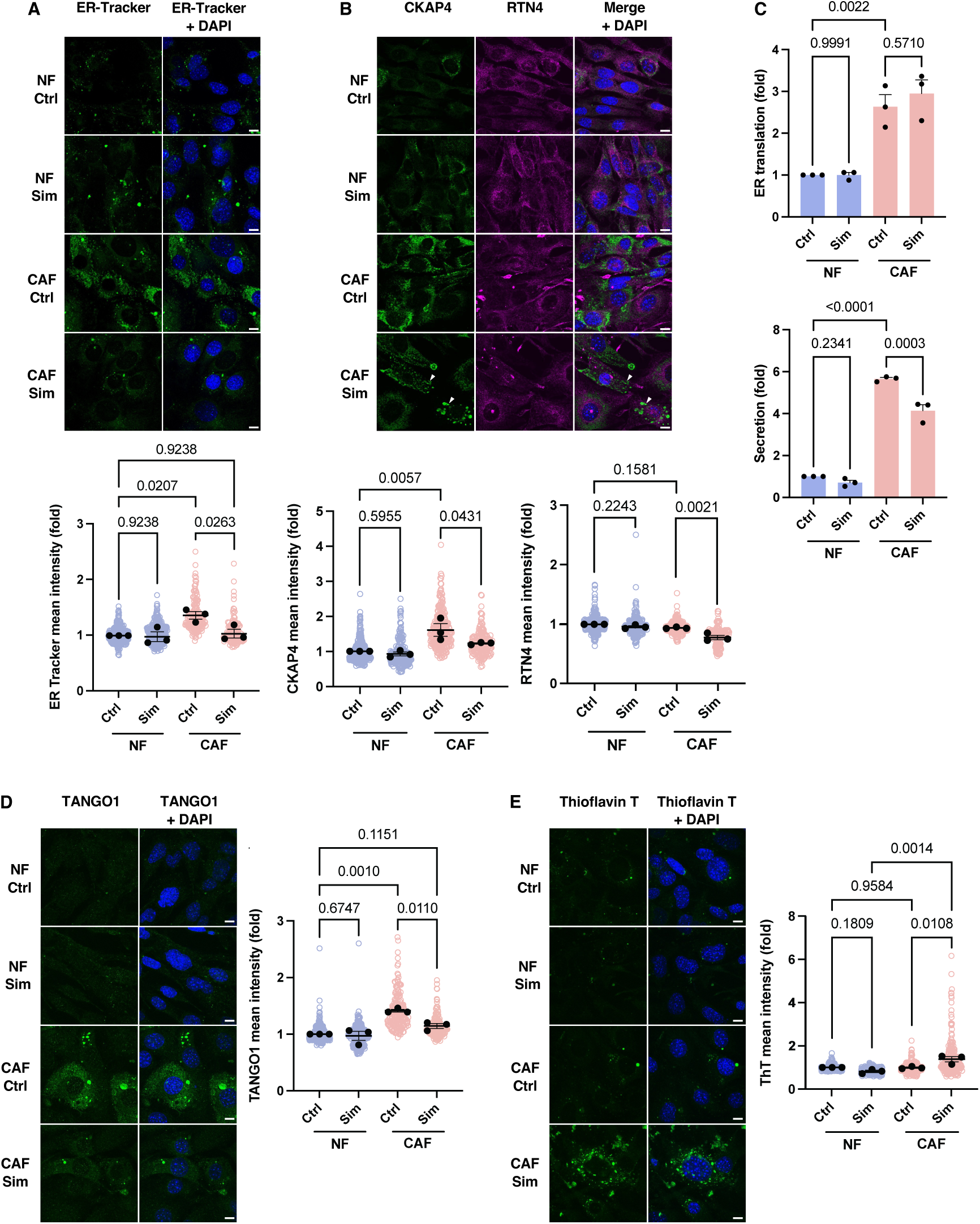
CAF ER function is dependent on cholesterol biosynthesis. **(A)** Representative ER-Tracker staining (green) in NFs and CAFs treated with DMSO (Ctrl) or 1 µM simvastatin (Sim) for 48 hours, and quantification of ER-Tracker mean intensity per cell, n=3 independent experiments (NF Ctrl=371 cells, NF Sim=354 cells, CAF Ctrl=168 cells, CAF Sim=138 cells). Nuclei are stained with DAPI (blue). Scale bars represent 10 µm. **(B)** Representative immunofluorescent staining of ER sheet marker CKAP4 (green) and ER tubule marker RTN4 (magenta) in NFs and CAFs treated with DMSO (Ctrl) or 1 µM Sim for 48 hours, and quantification of CKAP4 and RTN4 mean intensity per cell, n=3 independent experiments (NF Ctrl=345 cells, NF Sim=218 cells, CAF Ctrl=260 cells, CAF Sim=167 cells). Nuclei are stained with DAPI (blue). White arrows indicate disrupted CKAP4-stained sheet ER structures. Scale bars represent 10 µm. **(C)** Relative ER translation capacity (intracellular EGFP fluorescence intensity) and relative ER secretory capacity (extracellular GLuc activity) in GLuc-T2A-EGFP-expressing NFs and CAFs treated with DMSO (Ctrl) or 1 µM Sim for 48 hours, n=3. **(D)** Representative IF staining of TANGO1 (green) in NFs and CAFs treated with DMSO (Ctrl) or 1 µM Sim for 48 hours, and quantification of TANGO1 mean intensity per cell, n=3 independent experiments (NF Ctrl=422 cells, NF Sim=352 cells, CAF Ctrl=193 cells, CAF Sim=171 cells). Nuclei are stained with DAPI (blue), scale bars represent 10 µm. **(E)** Representative ThT staining (green) in NFs and CAFs treated with DMSO (Ctrl) or 1 µM Sim for 48 hours, and quantification of ThT mean intensity per cell, n=3 independent experiments (NF Ctrl=258 cells, NF Sim=154 cells, CAF Ctrl=184 cells, CAF Sim=230 cells). Nuclei are stained with DAPI (blue). Scale bars represent 10 µm.

### CAF ECM-remodelling function is dependent on cholesterol biosynthesis

Having determined that inhibition of cholesterol biosynthesis impairs ER sheet function, including the formation of TANGO1-positive ERES required for the export of ECM proteins, we next interrogated the role of cholesterol in regulating the ECM-remodelling capacity of CAFs. RNA-Seq analysis revealed downregulation of a range of ECM-related genes, and other myCAF markers, in CAFs following simvastatin treatment (**Supplementary Figure 4A,B,C**). Indeed, the gene encoding collagen XII (*Col12a1*), a fibril-associated collagen with interrupted triple helix (FACIT) collagen associated with metastasis (35), was the most significantly downregulated transcript in CAFs exposed to simvastatin (**Supplementary Figure 4A**). In alignment with the observed transcriptional response, collagen XII was downregulated at the protein level upon pharmacological inhibition of HMGCR (**Supplementary Figure 4D**). Consistent with our finding that inhibition of cholesterol biosynthesis suppresses ER function and induces protein misfolding in CAFs, upregulation of genes associated with the ER stress response and UPR was observed in CAFs following simvastatin treatment (**Supplementary Figure 4B**). Remarkably, 223 out of the 266 transcripts downregulated in CAFs following simvastatin treatment were also upregulated in CAFs relative to NFs (**Supplementary Figure 4E**), suggesting that inhibition of cholesterol biosynthesis contributes to normalisation of the myCAF phenotype. Critically, collagen deposition and collagen remodelling capacity were reduced in simvastatin-treated and HMGCR-depleted CAFs (**Figure 5A-D**). Collectively, these findings indicate that increased cholesterol biosynthesis is a metabolic adaptation that may be targeted in CAFs to abrogate ECM secretion and ECM remodelling capacity.

**Figure 5:**
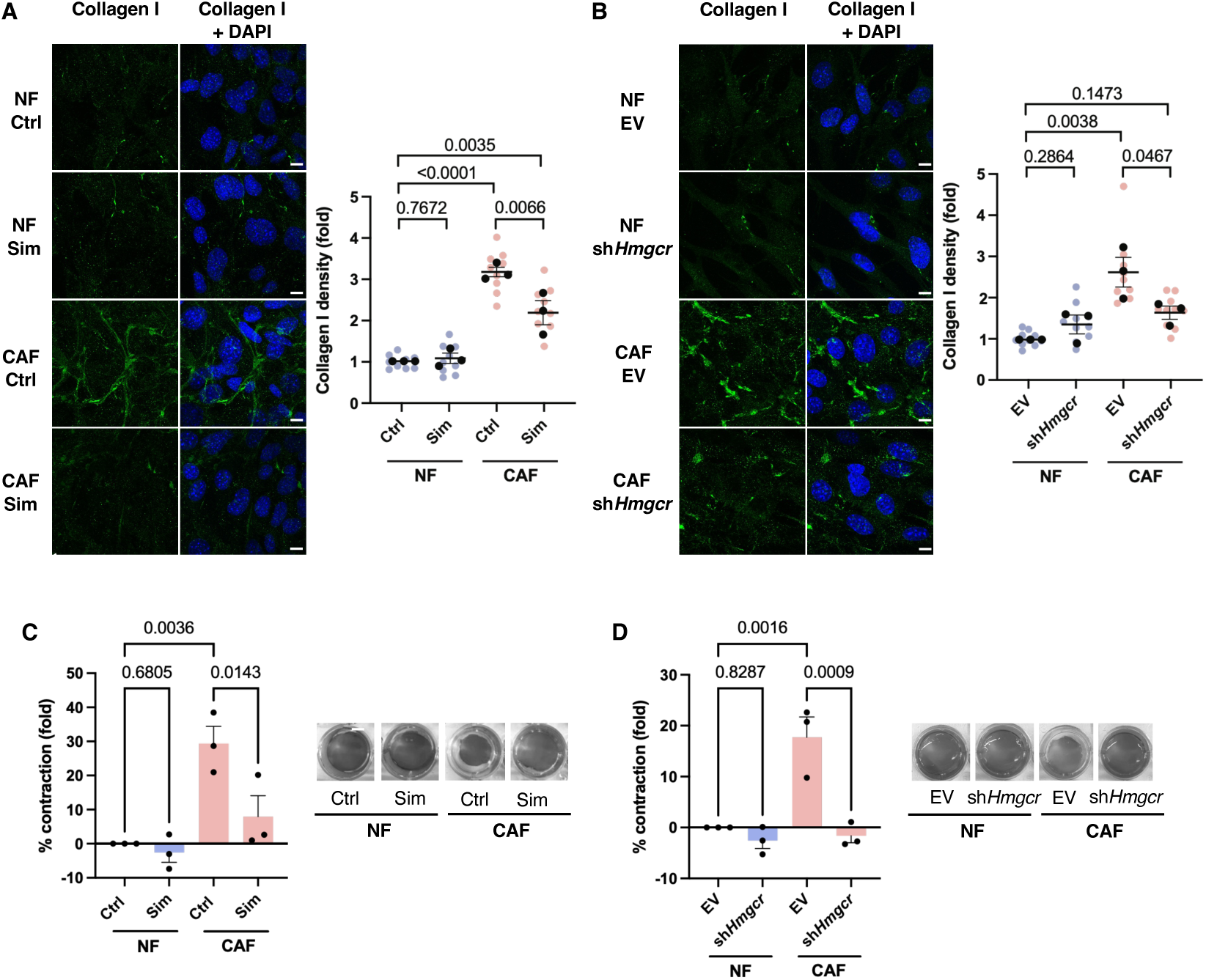
CAF ECM-remodelling function is dependent on cholesterol biosynthesis. Representative IF staining of collagen I (green) in **(A)** NFs and CAFs treated with DMSO (Ctrl) or 1 µM Sim for 48 hours, or **(B)** NFs and CAFs transduced with empty vector (EV) or sh*Hmgcr* constructs, and quantification of collagen I high-density matrix area normalized to cell number. Nuclei are stained with DAPI (blue). Scale bars represent 10 µm. ECM remodelling capacity of **(C)** NFs and CAFs treated with DMSO (Ctrl) or 1 µM Sim for 48 hours, or **(D)** NFs and CAFs transduced with EV or sh*Hmgcr* constructs as determined by a collagen contraction assay over 24 hours. Data are represented as % contraction relative to NF Ctrl **(C)** or NF EV **(D)**, n=3.

### Inhibition of cholesterol biosynthesis in CAFs reduces cancer cell invasion and metastasis

CAF-mediated reorganization of the breast tumour ECM is a potent driver of cancer cell invasion and metastasis (20,35). We thus sought to determine whether inhibition of cholesterol biosynthesis suppresses the ability of CAFs to promote tumour progression. Importantly, genetic depletion of *Hmgcr* effectively suppressed the ability of CAFs to remodel collagen plugs to permit invasion of PyMT cancer cells (**Figure 6A**). Having established a role for cholesterol biosynthesis in promoting pro-invasive ECM remodelling by CAFs *in vitro*, we turned to validate these findings in the *in vivo* context. To this end, wild-type or *Hmgcr*-deficient CAFs were co-implanted with PyMT cancer cells into the 4^th^ mammary fat pad of FVB/n mice (**Figure 6B**). Although no change in tumour latency, tumour weight, or survival of tumour-bearing mice was observed (**Supplementary Figure 5A-D**), *Hmgcr* depletion led to a striking reduction in metastatic burden, which was primarily driven by a decrease in the number of metastases (**Figure 6C,D, Supplementary Figure 5E**), although a trend towards decreased size of metastatic lesions was also observed (**Supplementary Figure 5F**). Overall, these data provide compelling evidence that reprogramming of cholesterol biosynthesis promotes the pro-tumorigenic function of myCAFs and facilitates cancer cell invasion and metastasis.

**Figure 6:**
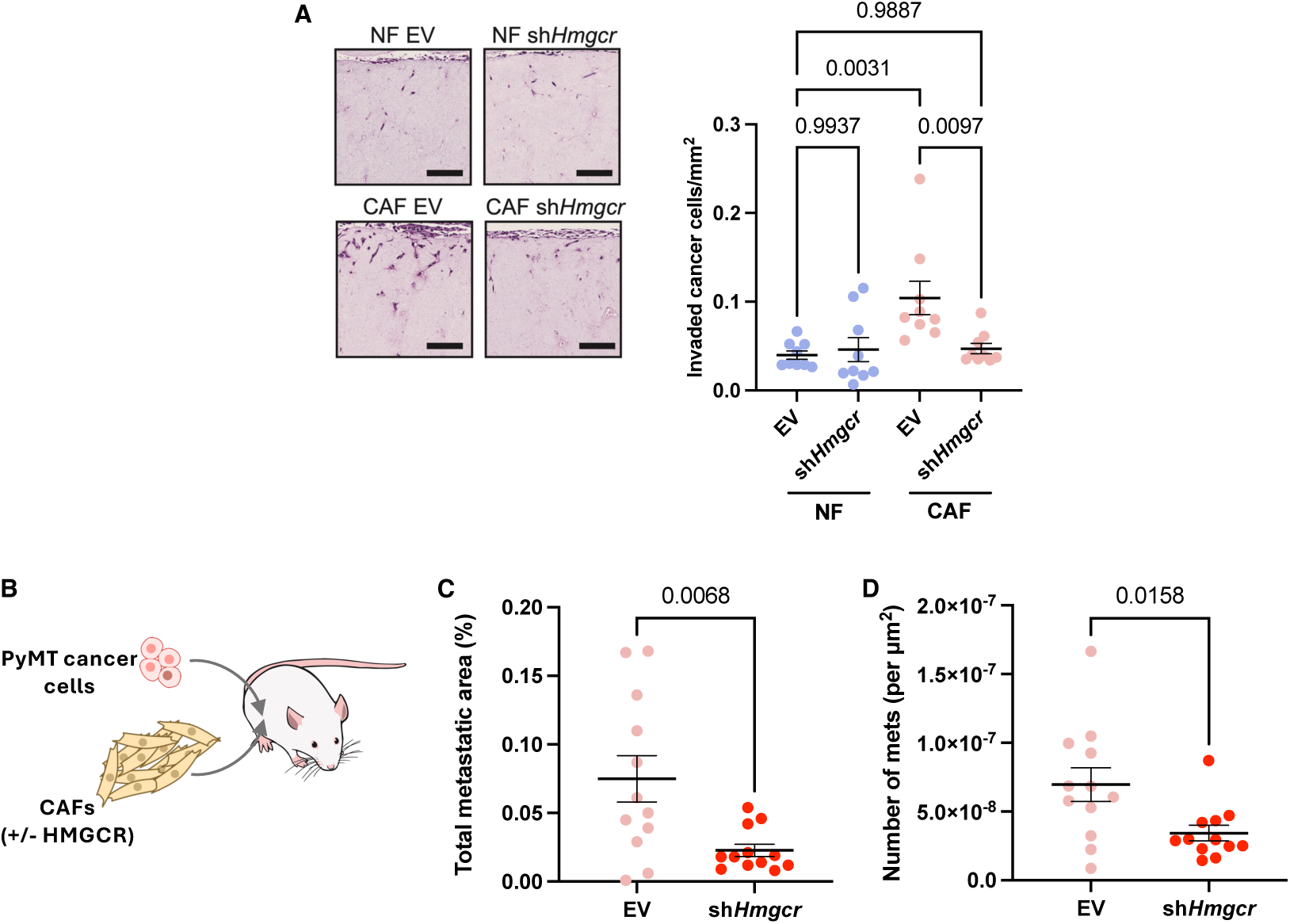
Inhibition of cholesterol biosynthesis in CAFs reduces cancer cell invasion and metastasis. **(A)** Representative H&E staining (left) and quantification (right) of PyMT cancer cell invasion into organotypic collagen matrices remodelled by NFs and CAFs transduced with empty vector (EV) or sh*Hmgcr* constructs, n=9 technical replicates per group from 3 independent experiments. **(B)** Schematic diagram of orthotopic co-implantation of PyMT cancer cells and CAFs. **(C)** Quantification of metastatic burden in mice co-injected with PyMT cancer cells and CAFs transduced with either EV or sh*Hmgcr* constructs, n= 12 mice per group. **(D)** Quantification of the number of lung metastases in mice co-injected with PyMT cancer cells and CAFs transduced with either EV or sh*Hmgcr* constructs, n= 12 mice per group.

## Discussion

Metabolic reprogramming is a hallmark of many cell types within the tumour microenvironment. Indeed, emerging evidence suggests that rewiring of cellular metabolism and nutrient acquisition strategies is fundamental to the pro-tumorigenic functions of CAFs. Such studies have largely emphasised the reprogramming of pathways that supply amino acids required for collagen biosynthesis (21,36). In this study, we reveal that reprogramming of cholesterol biosynthesis in CAFs indirectly supports ECM production and ECM remodelling by supporting ER function. Specifically, we show that cholesterol biosynthesis is upregulated in CAFs, and that cholesterol is enriched within the ER in these cells. Importantly, we find that ER sheet mass and secretory function are increased in CAFs and demonstrate that these phenomena are dependent on cholesterol biosynthesis and selective upregulation of the XBP1-mediated arm of the UPR. Finally, we find that inhibition of cholesterol biosynthesis suppresses collagen secretion and ECM-remodelling capacity *in vitro*, thus normalising CAF function. This cholesterol-dependent regulation of CAF function likewise modulates tumour progression *in vivo*, with CAF-specific disruption of cholesterol biosynthesis impairing the metastatic potential of cancer cells. Together, these findings implicate cholesterol biosynthesis as a critical regulator of the pro-tumorigenic activity of CAFs.

The ER is a highly dynamic organelle that plays numerous roles in the regulation of cellular metabolism, signalling and homeostasis. It is thus unsurprising that remodelling of ER architecture appears to be intrinsically linked with the establishment of unique cellular phenotypes. For instance, cells that secrete large amounts of protein such as pancreatic acinar cells and intestinal goblet cells contain abundant sheet ER, whilst highly lipogenic and steroidogenic cells such as adipocytes and testicular Leydig cells contain high levels of tubular ER (37–39). The differentiation of B lymphocytes into antibody-secreting plasma cells is likewise associated with a dramatic, XBP1-dependent increase in ER volume (40–42). Critically, the ER hosts several processes required for ECM synthesis, secretion and assembly (43). For example, procollagen molecules are translated, assembled into triple helices, and subjected to post-translational modification (PTM) via hydroxylation and glycosylation in the ER, before being secreted into the extracellular space via the ER-to-Golgi intermediate compartment (ERGIC) (44). Indeed, multiple ER-localised processes, such as HSP47-mediated triple helix stabilization and TANGO1-mediated expansion of coat protein complex II (COPII)-dependent vesicles, have evolved specifically to facilitate the correct folding and secretion of large collagen molecules (29,45,46). In this study, we demonstrated that CAFs exhibit a robust XBP1-dependent expansion of ER mass relative to their normal counterparts, as well as an increase in associated protein-folding and secretory function. This finding is particularly significant given that aberrant UPR signalling has been previously implicated in the aetiology of fibrotic pathologies (47–49). Our work suggests that remodelling of the ER is a critical adaptation underpinning the desmoplasia-promoting myCAF phenotype.

Cholesterol typically constitutes only 3-6% of the total lipid composition of the ER membrane, compared with 30-50% of the plasma membrane (50). However, our finding that ER secretory function is dependent on cholesterol synthesis is consistent with multiple previous studies. For instance, inhibition of cholesterol synthesis results in impaired lateral mobility of secretory membrane proteins in the ER, resulting in a failure of these proteins to reach ER exit sites (ERES) and subsequently be transported to the Golgi body (33,34). Moreover, depletion of cholesterol results in reduced turnover of COPII components at ERESs, suggesting that cholesterol is required for COPII-dependent carrier formation (34). Notably, ERESs have also been found to be particularly enriched for cholesterol compared with other regions of the ER membrane (51). Our study thus supports the notion that cholesterol supports ER secretory function by facilitating the formation of competent ERESs.

Whilst we have identified that reprogramming of cholesterol metabolism sustains CAF function in a cell-autonomous manner, multiple lines of evidence indicate that CAFs also supply lipid species to adjacent tumour cells. For instance, recent data suggest that lipid droplet-laden CAFs in *SETD2*-deficient pancreatic tumours sustain oxidative phosphorylation (OXPHOS) and tumour growth by transferring lipids to cancer cells (52). Moreover, uptake of CAF-derived fatty acids by colorectal cancer cells has been shown to promote migration (53). Our findings demonstrating upregulation of enzymes involved in cholesterol biosynthesis in MMTV-PyMT mammary CAFs corroborate previous analysis of patient-derived breast tumour stroma, which indicates that stromal expression of squalene epoxidase (*SQLE*) is associated with poor clinical prognosis (26). Moreover, the identification of cholesterol biosynthesis as a metabolic vulnerability in CAFs is particularly significant given previous studies demonstrating the anti-fibrotic properties of itraconazole; an anti-fungal agent that inhibits the cholesterol biosynthetic enzyme, lanosterol 14a-demethylase (CYP51A1) (54–56). Although these studies have attributed the CAF-inhibitory nature of itraconazole to its ability to suppress hedgehog (Hh), vascular endothelial growth factor (VEGF) and mitogen-activated protein kinase (MAPK) signalling, our findings pose an alternate mechanism through which itraconazole may target CAF function.

A range of studies have alluded to the anti-cancer properties of statins. For example, statin use is associated with lowered incidence of several cancers, as well as more favourable prognosis and reduced rates of recurrence and mortality (57–61). The anti-tumorigenic effects of statins have primarily been attributed to the disruption of cancer cell-intrinsic processes, such as lipid raft-mediated signalling and the isoprenylation of oncogenic proteins such as RAS and Rho (62–64). Our findings provide an additional mechanism by which statins can target tumour progression at the microenvironmental level. This is particularly pertinent as the treatment of CAF-driven desmoplasia remains an area of unmet clinical need. Our work also has important potential implications for other conditions in which myofibroblasts produce pathological fibrosis, such as metabolic dysfunction-associated steatohepatitis (MASH) and idiopathic pulmonary fibrosis. Indeed, treatment with atorvastatin has been shown to attenuate liver fibrosis in a mouse model of MASH (65). Given that statins are already widely used and well-tolerated, repurposing them as an anti-fibrotic agent may hold significant therapeutic potential.

In conclusion, this study advances our understanding of the fundamental biology of myCAFs, identifying cholesterol-dependent remodelling of the ER as an adaptive process that sustains collagen secretion and pro-tumorigenic ECM-remodelling capacity. Our work provides impetus to further explore cholesterol biosynthesis as a metabolic vulnerability in CAFs that may be targeted to ameliorate tumour desmoplasia.

## Methods

### Cell lines and cell culture

293T cells (ATCC; CRL-3216) were cultured in DMEM, high glucose supplemented with 10% heat-inactivated FBS and 1 mM sodium pyruvate. Mammary fibroblasts derived from wild-type FVB/n mice (NFs), or cancer-associated fibroblasts from transgenic PyMT FVB/n mice (CAFs) were a kind gift from Fernando Calvo (IBBTEC, Spain) (22). NFs and CAFs were maintained in DMEM, high glucose supplemented with 10% FBS, 1 mM sodium pyruvate and 1% Insulin-Transferrin-Selenium (ITS-G). PyMT 20065 cancer cells were derived with the support of Karen Blyth at the CRUK Beatson Institute in Glasgow through the SEARCHbreast initiative (https://searchbreast.org/). PyMT 20065 cancer cells were maintained in DMEM, high glucose supplemented with 10% FBS 1% penicillin/streptomycin, 5 μg/mL insulin, 10 ng/mL epidermal growth factor and 10 ng/mL Cholera Toxin. Plasmax was prepared in-house as previously described (23). 72 hours prior to the experimental period, NFs and CAFs were transitioned to Plasmax supplemented with 10% FBS and 1% ITS-G and Plasmax media was refreshed every 48 hours. Cells were maintained in a humidified incubator at 37°C with 5% CO_2_ and regularly assayed for mycoplasma contamination.

### Generation of stable cell lines

sh*Xbp1* (TRCN0000232021) and sh*Hmgcr* (TRCN0000173307) shRNA sequences were derived from The RNAi Consortium (TRC) MISSION shRNA Library and are listed in Table 1. All shRNAs were cloned into pLKO.1 hygro (a gift from Bob Weinberg; Addgene plasmid #24150, RRID:Addgene_24150). shRNA knockdown and GLuc-T2A-EGFP-expressing lines were generated through lentiviral transduction. Lentiviral particles were produced by co-transfecting 293T cells with pLKO.1 hygro lentiviral constructs or pLV GLuc-T2A-EGFP (VectorBuilder), along with pMDLg/pRRE, pRSV-Rev, and pMD2.g (gifts from Didier Trono. Addgene plasmid #12251, RRID:Addgene_12251; Addgene plasmid #12253, RRID:Addgene_12253; Addgene plasmid #12259, RRID:Addgene_12259). Target cells were transduced using 0.45μm-filtered viral supernatant containing 10 μg/mL polybrene. Cells were maintained in a humidified incubator at 37°C with 5% CO_2_ for 48 hours before selection. NFs and CAFs transduced with pLKO.1 hygro shRNA constructs were selected with 1 mg/mL hygromycin for 48 hours, while NFs and CAFs transduced with pLV-GLuc-T2A-EGFP were selected with 20 μg/mL blasticidin for 72 hours. Following complete selection, cells were maintained in selection antibiotic at 50% of the initial selection concentration. Antibiotic was removed when cells were seeded for experiments.

### Simvastatin treatment

Cells were seeded in Plasmax supplemented with 10% FBS and 1% ITS-G. 24 hours after seeding, media was replaced with fresh media containing 1 μM simvastatin (Selleck Chemicals; S1792) prepared in dimethylsulfoxide (DMSO) (EMD Millipore; 317275). Controls were treated with vehicle (DMSO) alone. The duration of simvastatin treatment is specified in the relevant figure legend for each experiment.

### RNA sequencing (RNA-Seq) and analysis

Experimental conditions for RNA-Seq experiments are described in the relevant figure legends. Briefly, RNA was extracted using the NucleoSpin RNA isolation kit (Machery-Nagel) according to manufacturer’s guidelines. Libraries were prepared using the QuantSeq 3’ mRNA-seq kit (Lexogen) and sequenced with an Illumina NextSeq 500, with single-end 75 bp reads to a depth of 5-15 million reads per sample. Raw read quality was assessed using FastQC. FastQ files were trimmed using Cutadapt (v2.1) and mapped to the mouse reference genome (Ensembl mm10 GRCm38) using HISAT2 (v2.2.0). Samtools (v1.9) was used to convert SAM files to BAM format and to sort BAM files. Counts were generated using featureCounts (v1.6.0). Filtering and normalisation of data was carried out using edgeR (v4.6.3) (66) and differential gene expression analysis was carried out using limma-voom (v3.64.3) (67). Gene set enrichment analysis (GSEA) was carried out with clusterProfiler (v4.16.0) (68) using the “gseGO” function (biological process (“BP”) and cellular component (“CC”) ontologies). GSEA plots were generated with enrichplot (v1.28.4) (69) using the “gseaplot2” function. Heatmaps were generated using pheatmap (v1.0.13) (70). Volcano plots were generated using VolcanNoseR (71). UpSet plots were generated using UpSetR (v1.4.0) (72).

### Analysis of Human Breast Cancer Atlas single cell RNAseq (scRNAseq) data

Patient-derived breast cancer scRNAseq data (25) (GSE176078) were accessed via the Single Cell

Portal (https://singlecell.broadinstitute.org/single_cell). Briefly, this data was derived from analysis of 26 human primary breast cancer tumours via scRNAseq using the Chromium platform (10× Genomics). Data were filtered to show stromal cell clusters. Gene expression is represented by log-normalised expression values.

### Quantitative polymerase chain reaction (qPCR)

RNA was extracted using the NucleoSpin RNA isolation kit (Machery-Nagel) according to manufacturer’s guidelines. 0.5 – 1 μg of RNA was reverse-transcribed to complementary DNA (cDNA) using the High-Capacity cDNA Reverse Transcription Kit (Thermo Fisher Scientific). cDNA was diluted 5-fold in nuclease-free H_2_O. Each qPCR reaction contained 4 μL diluted cDNA, 0.5 μL nuclease-free H_2_O, 1.5 μL 2 μM forward and reverse primers and 7.5 μL Fast SYBR™ Green Master Mix (Thermo Fisher Scientific). qPCR was run in MicroAmp Fast Optical 96- or 384-Well Reaction Plates (Thermo Fisher Scientific) using a CFX Touch Real-Time PCR Detection System (Bio-Rad). Relative gene expression was analyzed using the ΔΔCt method (73) normalized to *Rplp0.* qPCR primer sequences are listed in Table 2.

### Immunoblotting

Cells at 80% confluency were washed with cold PBS immediately prior to lysis on ice with cold RIPA buffer (1% NP-40, 0.5% sodium deoxycholate, 0.1% SDS, 150 mM NaCl, 50 mM Tris-HCl, pH 7.5) containing 1X protease inhibitor cocktail (Sigma-Aldrich) and 1X phosphatase inhibitor (PhosSTOP; Sigma-Aldrich). Lysates were collected by scraping and debris was pelleted by centrifugation at 16,000 x *g* for 10 minutes at 4°C. Protein concentration was determined by Detergent-Compatible (DC) assay (Bio-Rad) as per manufacturer’s instructions. Protein samples were resolved using either 15% Tris-glycine gels (made in-house) or NuPAGE™ 4-12% Bis-Tris gels (Thermo Fisher Scientific; NP0336) (Table 3). For Tris-glycine SDS-PAGE, equal quantities of protein lysate were combined with 5X Laemmli sample loading buffer (312.5 mM Tris-HCl (pH 6.8), 25% glycerol, 10% SDS, bromophenol blue) containing 5% β-mercaptoethanol and incubated at 90°C for 10 minutes. Samples were resolved in SDS-PAGE running buffer (25 mM Tris, 250 mM glycine, 0.1% SDS) using a Mini-PROTEAN Tetra Vertical Electrophoresis Cell (Bio-Rad; 1658004). Resolved proteins were transferred to nitrocellulose membrane at 100V for 60-75 minutes in transfer buffer (25 mM Tris, 192 mM glycine, 20% methanol) using a Mini Trans-Blot® System (Bio-Rad). For Bis-Tris SDS-PAGE, equal quantities of protein were suspended in NuPAGE™ LDS sample buffer (Thermo Fisher Scientific) containing 1X NuPAGE™ sample reducing agent (Thermo Fisher Scientific; NP0009). Samples were incubated at 70°C for 10 minutes. Samples were resolved on NuPAGE™ 4-12% Bis-Tris protein gels in a Mini Gel Tank (Thermo Fisher Scientific) using either NuPAGE™ MOPS Running Buffer (Thermo Fisher Scientific) for high molecular weight targets, or NuPAGE™ MES Running Buffer (Thermo Fisher Scientific) for low to medium molecular weight targets. Resolved proteins were transferred to nitrocellulose membrane at 23V for 75 minutes at 4°C in a Mini Gel Tank using NuPAGE™ Transfer Buffer (Thermo Fisher Scientific). Membranes were blocked for an hour in TBS containing 2.5% bovine serum albumin (BSA). Membranes were washed once with TBS-T and 3 times with Milli-Q water. Blocked membranes were incubated overnight at 4°C with primary antibody diluted in TBS containing 5% BSA. Membranes were washed and incubated with IRDye secondary antibodies (LI-COR Biosciences) diluted in TBS containing 2.5% BSA and imaged using the Odyssey CLx imaging system (LI-COR Biosciences). Antibodies and antibody dilutions are listed in Table 3.

### Glucose labelling

Glucose labelling was based on a previously described protocol (24). Briefly, cells were seeded in Plasmax supplemented with 10% FBS and 1% ITS-G at 0.1 x 10^6^ cells per well in duplicate wells in a 6 well plate. 24 hours after seeding, cells were spiked with 0.2 μCi [U-^14^C]glucose (PerkinElmer) and incubated for an additional 24 hours in a humidified incubator at 37°C with 5% CO_2_. Cells were washed twice with PBS before lysis in Trizol™ Reagent. Trizol lysates were fractionated according to manufacturer’s instructions to obtain polar, RNA, DNA, organic and protein fractions. The polar fraction was derived by retaining supernatant from RNA precipitation, while the organic fraction was derived by retaining supernatant from protein precipitation. Each fraction was combined with 4 mL Optiphase HiSafe 3 (PerkinElmer), quantified using a Tri-Carb 2910 TR Liquid Scintillation Analyzer (PerkinElmer) and normalized to cell number from duplicate wells. Total [U-^14^C]glucose uptake was determined by calculating the sum of ^14^C-labelling across all fractions.

### Lipogenesis assay

A previously described protocol was employed with minor modifications (74,75). Briefly, cells were seeded in Plasmax supplemented with 10% FBS and 1% ITS-G at 0.1 x 10^6^ cells per well in a 6 well plate and allowed to adhere for 24 hours before being spiked with 0.2 μCi [1-^14^C]acetate (PerkinElmer) for the final 4 hours of the experimental period. Cells were washed twice with PBS before lysis in 0.5% Triton X-100 (Sigma-Aldrich). Lipids were extracted with 2:1 (v/v) chloroform/methanol followed by centrifugation at 1000 rpm for 20 min. 150 μL of ^14^C-labelled lipid in the denser fraction was combined with 4 mL Optiphase HiSafe 3, quantified using a Tri-Carb 2910 TR Liquid Scintillation Analyzer and normalized to cell number.

### Cholesterol assay

Total cholesterol was quantified using a Cholesterol/Cholesteryl Ester Assay Kit (Abcam). Briefly, cells were seeded at 0.5 x 10^6^ cells per 6 cm plate in Plasmax supplemented with 10% FBS and 1% ITS-G. After 48 hours, cells were harvested, and 0.8 x 10^6^ cells were aliquoted per condition. Total cholesterol was extracted and quantified as per the manufacturer’s instructions (colorimetric protocol) using a Cytation™ 3 plate reader (BioTek).

### ER fractionation and cholesterol quantification

A previously described protocol (76,77) was adapted to quantify cholesterol in the ER-enriched subcellular fraction. Briefly, NFs and CAFs grown in Plasmax supplemented with 10% FBS and 1% ITS-G were harvested and counted. 15 x 10^6^ cells were washed with cold PBS and centrifuged at 300 x *g* for 4 minutes. The cell pellet was resuspended in 250 μL ice-cold STM Buffer (250 mM sucrose, 5 mM MgCl_2_, 50 mM Tris HCl pH 7.4) containing protease inhibitor cocktail (Sigma-Aldrich) and phosphatase inhibitor (Sigma-Aldrich) and homogenized with a motorized pestle for 5 minutes on ice before centrifugation at 800 x *g* for 15 minutes at 4°C. Supernatant (S/N I) was transferred to a new tube. The remaining pellet was resuspended in 1.25 mL ice-cold STM Buffer containing protease inhibitor cocktail and phosphatase inhibitor and homogenized with a motorized pestle for 10 minutes on ice, before centrifugation at 800 x *g* for 15 minutes at 4°C to pellet the nuclear fraction. Supernatant (S/N II) was transferred to a new tube. S/N I and S/N II were centrifuged in separate tubes at 6,000 x *g* for 15 minutes at 4°C to pellet the mitochondrial fraction. Remaining supernatants were combined and transferred to duplicate ultracentrifuge tubes (Beckman-Coulter, 344059). Samples were ultracentrifuged at 100,000 x *g* for 1 hour using a Beckman Coulter XPN-100 ultracentrifuge equipped with a SW 41 Ti swinging bucket rotor (Beckman-Coulter) to pellet the microsomal (ER-enriched) fraction. One replicate microsomal pellet for each cell type was resuspended in SDS lysis buffer (1% SDS, 50 mM Tris-HCl, 10 mM EDTA) containing protease inhibitor, phosphatase inhibitor and Pierce™ Universal Nuclease (Thermo Fisher Scientific) for protein quantification via DC assay and immunoblotting. The remaining microsomal pellet was resuspended in 200 μL methanol/water (80:20). 50 μL of this solution was extracted with 125 μL butanol/methanol (1:1) by vortexing followed by centrifugation for 15 minutes at 4,000 x *g* at 4°C. Supernatant was collected and vacuum dried. Cholesterol was quantified using the Cholesterol/Cholesteryl Ester Assay Kit (Abcam) as per manufacturer’s instructions (fluorometric protocol) and normalized to microsomal protein. Fractionation was validated by immunoblotting.

### Gaussia luciferase assay

NFs and CAFs stably expressing GLuc-T2A-EGFP were seeded in Plasmax supplemented with 10% FBS and 1% ITS-G at 0.1 x 10^6^ cells per well in a 6 well plate (0.05 x 10^6^ cells for simvastatin experiments). 24 hours before the experimental end point, media was replaced. A 50 μL aliquot of culture media was collected to quantify baseline GLuc. 24 hours later, a 50 μL aliquot of culture media was collected to quantify GLuc secretion. Cells were trypsinized and counted. 0.1 x 10^6^ cells were resuspended in PBS per condition for quantification of intracellular EGFP. Secreted GLuc in cell culture media was quantified with the Pierce™ Gaussia Luciferase Glow Kit (Thermo Fisher Scientific) according to the manufacturer’s instructions. EGFP fluorescence intensity and luminescence were detected using a Cytation™ 3 plate reader. Luminescence detected in baseline samples was subtracted from that detected at experimental endpoint. GLuc secretion was determined by normalizing luminescence to total cell count for each condition.

### Filipin staining

Cells were seeded in Plasmax supplemented with 10% FBS and 1% ITS-G on poly-L-lysine-coated 8 well chamber slides (Ibidi; 80826) at 0.02 x 10^6^ cells per chamber and maintained in a humidified incubator at 37°C with 5% CO_2_ for 48 hours. Media was removed and cells were washed twice with PBS before fixation with freshly prepared 4% PFA for 15 minutes at room temperature. Fixed cells were washed with PBS and stained with freshly prepared 50 μg/mL Filipin III (Sigma-Aldrich) in PBS for 1 hour at room temperature in the dark. Filipin was removed, and cells were washed and imaged in PBS containing 5 μM Draq5 staining solution (Miltenyi Biotec) using a Zeiss Elyra PS.1 confocal microscope (Zeiss) equipped with a 63X/1.4 oil DIC objective.

### Immunofluorescent (IF) staining

Cells were seeded in Plasmax supplemented with 10% FBS and 1% ITS-G on poly-L-lysine-coated 8 well chamber slides (Ibidi; 80826) at 0.02 x 10^6^ cells per chamber. After 24 hours, media was replaced with Plasmax containing 10% FBS, 1% ITS-G and 5 μg/mL L-ascorbic acid. Fresh L-ascorbic acid was added every 24 hours for a total of 72 hours, or for the duration of the experimental period where otherwise indicated. Cells were washed with PBS and fixed for 10 minutes at room temperature using freshly prepared 4% PFA in PBS. Fixed cells were washed twice with PBS. Permeabilization was carried out for intracellular staining by incubating fixed cells in cold 70% ethanol in PBS for 30 minutes on ice. Cells were washed with PBS and blocked in IF Blocking Buffer (1% BSA, 138 mM NaCl, 20 mM Tris-HCl pH 7.5) for 20 minutes at room temperature. Cells were washed with PBS and incubated in primary antibody diluted in IF blocking buffer for 2 hours at room temperature (antibody dilutions are specified in Table 3). Cells were washed with PBS and incubated in appropriate fluorescent secondary antibody diluted in IF blocking buffer for 2 hours at room temperature in the dark (antibody dilutions are specified in Table 3). Cells were washed with PBS three times and nuclei were stained with 0.5 μg/mL DAPI (Sigma-Aldrich) for 5 minutes at room temperature in the dark. Cells were washed twice with PBS and stored in DAKO fluorescence mounting medium (Agilent Technologies; S3023) at 4°C until imaging using an Olympus FV3000 Scanning Confocal Microscope (Olympus Life Science) equipped with silicone oil immersion 60X/1.42 objective. Mean fluorescence intensity of ER markers was quantified using CellProfiler (78). Briefly, nuclei were segmented using the IdentifyPrimaryObjects module and used as seed objects for segmentation of the cell body and cytoplasm using the IdentifySecondaryObjects and IdentifyTertiaryObjects modules. Intensity was quantified using the MeasureObjectIntensity module. High density matrix was quantified in FIJI using TWOMBLI (79) and normalised to cell number for each field of view.

### ER Tracker staining

Cells were seeded and maintained as described in “Immunofluorescent (IF) staining”. At the end of the experimental period, media was removed, and cells were washed with serum-free EBSS (Thermo Fisher Scientific). Cells were stained using pre-warmed 1 μM ER-Tracker™ Green (Thermo Fisher Scientific) in serum-free EBSS for 20 minutes in a humidified incubator at 37°C with 5% CO_2_. Staining solution was removed, and cells were equilibrated in complete Plasmax containing appropriate supplements for 10 minutes in a humidified incubator at 37°C with 5% CO_2_. Media was removed and cells were washed with PBS. Cells were fixed in 4% PFA for 2 minutes at 37°C. Fixed cells were washed with PBS, stained with DAPI (see “Immunofluorescent (IF) staining”) and stored in DAKO fluorescence mounting medium at 4°C until imaging. Cells were imaged and analyzed using the same method described in “Immunofluorescent (IF) staining”.

### Thioflavin T staining

Cells were seeded and maintained as described in “Immunofluorescent (IF) staining”. A 5 mM (1000X) stock of Thioflavin T (ThT, Sigma-Aldrich) was prepared fresh for each experiment by dissolving 8 mg ThT in ethanol. Cells were live stained with ThT in cell culture media at a final concentration of 5 μM for 40 minutes in a humidified incubator at 37°C with 5% CO_2_. Media was removed and cells were washed with PBS. Cells were fixed in 4% PFA for 20 minutes at room temperature. Fixed cells were washed with PBS, stained with DAPI (see “Immunofluorescence (IF) staining”) and stored in DAKO fluorescence mounting medium at 4°C until imaging. Cells were imaged and analyzed using the same method described in “Immunofluorescent (IF) staining”.

### Organotypic collagen remodelling assays

24 hour collagen remodelling assays were conducted as per previously described protocols (80,81). Briefly, NFs and CAFs cultured in Plasmax supplemented with 10% FBS and 1% ITS-G were harvested, counted, and 0.16 x 10^6^ cells were aliquoted in a microcentrifuge tube for each condition and brought to a total volume of 800 μL with culture media. Cell suspension was combined with 400 μL 3 mg/mL rat tail collagen I solution (Thermo Fisher Scientific) and immediately neutralized with 12 μL of sterile 10 mM NaOH solution. The cell-collagen solution was mixed and 500 μL was transferred into duplicate wells in a 24 well plate. Plates were incubated for 20 minutes in a humidified incubator at 37°C with 5% CO_2_, allowing collagen plugs to set. A pipette tip was used to detach edges of the collagen plug from the well and 600 μL fresh Plasmax supplemented with 10% FBS and 1% ITS-G was added. Cell-populated collagen matrices were incubated in a humidified incubator at 37°C with 5% CO_2_ for 24 hours. Media was removed, matrices were washed twice with PBS and fixed for 30 minutes at room temperature with 4% PFA. Fixative was removed, and matrices were washed with H_2_O before staining with Eosin Y (Agilent Technologies) for 2 minutes. Collagen matrices were washed with H_2_O followed by 100% ethanol and allowed to dry before imaging using the ChemiDoc Touch Imaging System (Bio-Rad) in ‘silver stain’ mode. The area of each collagen plug was determined using FIJI and averaged between technical duplicates. 7 day collagen remodelling assays were conducted as previously described (82). Briefly, a total of 0.15 x 10^6^ NFs or CAFs were embedded in 1.25 mL neutralized rat tail collagen (1.5 mg/mL) in 12 well plates. 3D organotypic matrices were maintained in Plasmax media supplemented with 10% FBS and 1% ITS-G. Matrices were allowed to contract over 7 days in a humidified incubator at 37°C with 5% CO_2_. Plates were imaged daily on a CCD flatbed scanner.

### Organotypic invasion assay

Following 7 days of contraction as described in “Organotypic collagen remodelling assays”, 0.05 x 10^6^ PyMT cancer cells were seeded on top of the remodelled collagen matrix and cultured in Plasmax media containing appropriate supplements over 2 days. Once cells had attached, matrices were carefully transferred to an air-liquid interface for cancer cell invasion. On day 7, organotypic matrices were formalin fixed and paraffin embedded for sectioning and H&E staining. Cell invasion was scored in three representative areas of each matrix and calculated as the number of invaded cells per mm^2^.

### Animal ethics

All animal procedures were conducted in accordance with the Australian Code of Practice for the Care and Use of Animals for Scientific Purposes by the National Health and Medical Research Council and were approved by the Animal Ethics Committee of the St Vincent’s Precinct and Garvan Institute (ARA 22_04). FVB/NJAusb female mice (7 weeks old) were purchased from Australian BioResources (ABR) and acclimatised for four weeks prior to experimentation. During acclimatisation, mice were monitored weekly by body weight measurements and general welfare inspections. Throughout the study, all animals were housed under a 12-hour light/dark cycle at standard ambient temperature and humidity, with ad libitum access to water and a standard diet (23% protein, 6% fat, 5% fibre).

### *In vivo* orthotopic co-implantation study

PyMT 20065 cancer cells (3 × 10⁵) were co-implanted with either EV or shHmgcr PyMT CAFs (9 × 10⁵; 1:3 ratio) into the fourth mammary fat pad of 11-week-old female mice (12 mice per group). Cells were resuspended as single-cell suspensions in 1X Hanks’ Balanced Salt Solution (HBSS; Gibco #14175-095) and kept on ice until injection (50 µL). All mice developed tumours. Mice were monitored daily during the first week post-implantation and three times weekly thereafter by body weight measurement and welfare inspection. Tumour growth was measured using callipers, and tumour volume was calculated as: volume = (maximum dimension² × minimum dimension) × 0.52. Mice were euthanised by CO₂ asphyxiation when tumours reached the primary ethical endpoint of 520 mm³. Tumour and lung tissue was collected, and either snap-frozen or formalin-fixed (10% neutral-buffered formalin (NBF; Australian Biostain #ANBFC)) for 24 h followed by histological processing. Lungs were inflated with 1% carboxymethylcellulose solution (MedChemExpress #HY-Y0703) prior to formalin fixation.

### Histopathology and H&E staining

Following 24 hours of fixation in 10% NBF, tissues were transferred to 70% ethanol, before being processed and embedded in paraffin. Sectioned tumour and lung tissues (4 µm) were deparaffinised and rehydrated through a series of graded ethanol washes. Tumour and lung tissues were stained with H&E to quantify metastasis using a Leica ST5010 Autostainer XL. Slides were imaged using a Hamamatsu NanoZoomer S210 (NDP.Scan v3.4 software) and analysed in QuPath (v0.5). Lung metastases were quantified across four lung lobes by measuring and calculating (i) the total number of metastatic foci, (ii) the number of foci per µm² of tissue, (iii) the total metastatic burden (%), normalised to the total cross-sectional area, and (iv) the average area of metastatic lesion (µm²).

### Statistics

Statistical analyses were performed with Prism 10 (GraphPad Software). Unpaired Student’s t-tests were used for comparison between two groups. One-way analysis of variance (ANOVA) with Holm-Sidak correction for multiple comparisons was used for comparison of more than two groups. Plots are represented as mean values with error bars representing standard error of the mean (SEM). Unless otherwise indicated, n refers to independent biological replicates. Other statistical details of experiments can be found in the figure legends.

## Acknowledgements

S.V. is supported by an Australian Government Research Training Program Scholarship and the Peter MacCallum Cancer Foundation. E.T.Y.M. is supported by an Australian Government Research Training Program Scholarship and Tour de Cure. N.L.Y. and J.L.C. are supported by NHMRC Grants (GNT2013881 and GNT2033065). T.R.C. is supported by an NHMRC Investigator Grants (GNT2033065). K.K.B. is supported by NHMRC Ideas Grants (GNT2004212 and GNT2012313) and a Victorian Cancer Agency Mid-Career Research Fellowship (MCRF17020). A.G.C. is supported by a National Health and Medical Research Council (NHMRC) Investigator Grant (GNT1176650), and an Australian Research Council Discovery Project Grant (DP200102693). We acknowledge support from the Peter MacCallum Cancer Centre Foundation and the Australian Cancer Research Foundation. We extend our thanks to the Peter MacCallum Cancer Centre Core Facilities and their staff who provided support for this work; namely the Molecular Genomics Core (RRID:SCR_025695), the Centre for Advanced Histology and Microscopy (RRID:SCR_025432), and Research Laboratory Support Services (RRID:SCR_025699). We acknowledge the facilities, as well as scientific and technical assistance, at Australian Bioresources (ABR) and the Garvan Histology Core Facility. Finally, we thank members of the Brown Laboratory (Peter MacCallum Cancer Centre) and Cox Laboratory (Peter MacCallum Cancer Centre) for helpful discussions.

## Author contributions

S.V. and K.K.B. designed research; S.V., N.L.Y., E.T.Y.M., J.L.C., J.A.R., R.A.G., and R.M.S. performed research; S.V., N.L.Y., E.T.Y.M., and J.L.C. analyzed data; A.G.C., T.R.C., and K.K.B. supervision; and S.V. and K.K.B. wrote the paper.

---

**Supplementary Figure 1.**
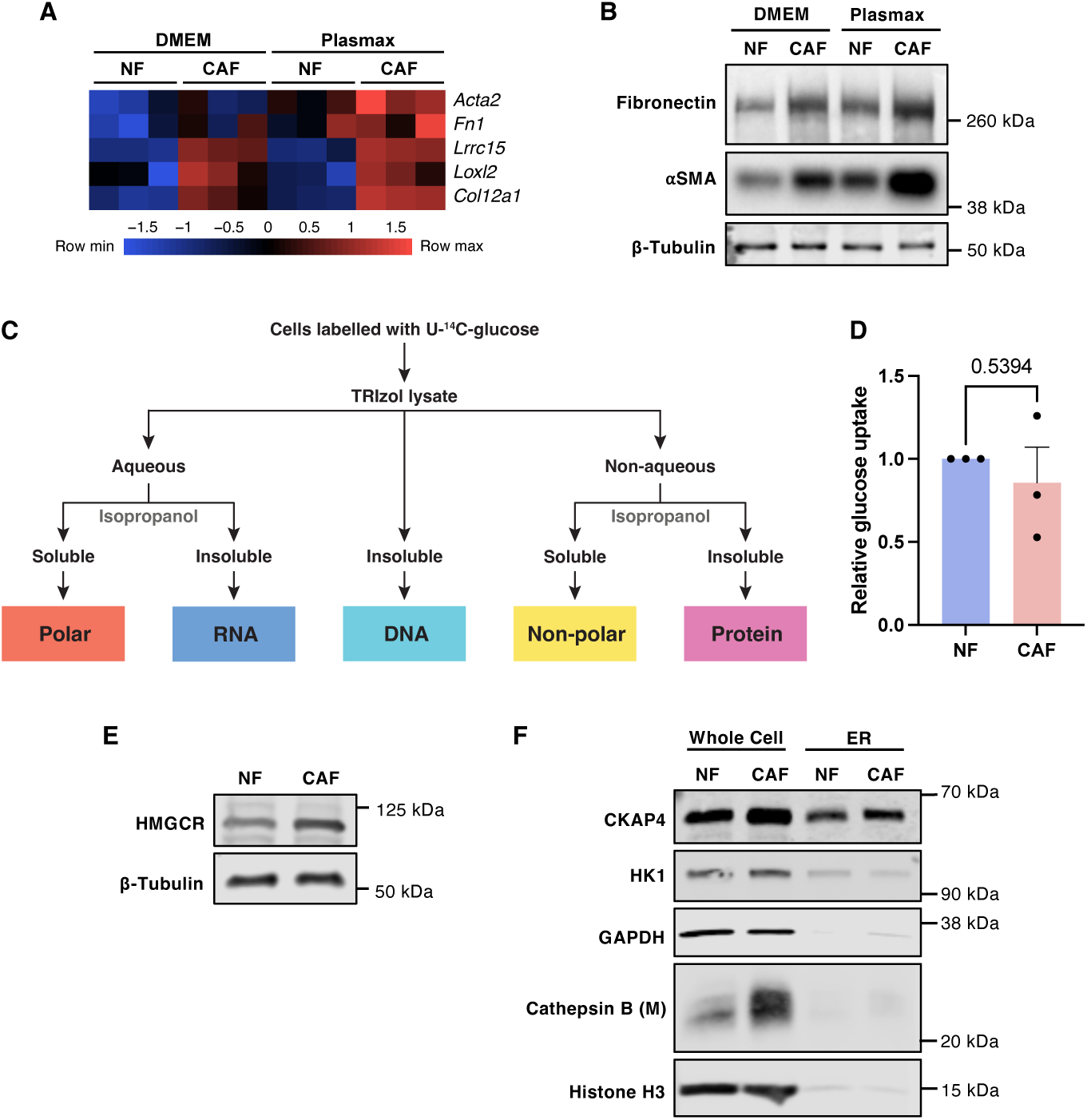
**(A)**  Heatmap of CAF marker gene expression in NFs and CAFs cultured in DMEM or Plasmax, as determined by RNA-seq analysis, n=3. **(B)** Representative immunoblot analysis of CAF marker expression in NFs and CAFs cultured in DMEM or Plasmax. β-tubulin serves as a loading control. **(C)** Schematic of approach used to monitor glucose incorporation into the major classes of biological macromolecules. **(D)** Relative [U-^14^C]glucose uptake in CAFs versus NFs, n=3. **(E)** Representative immunoblot analysis of HMGCR expression in NFs and CAFs. β-tubulin serves as a loading control. **(F)** Immunoblot analysis of whole cell and ER-enriched fractions derived from NFs and CAFs. CKAP4, HK1, GAPDH, Cathepsin B (M = mature) and Histone H3 serve as markers of ER, mitochondria, cytoplasm, lysosomes and nuclei respectively.

**Supplementary Figure 2.**
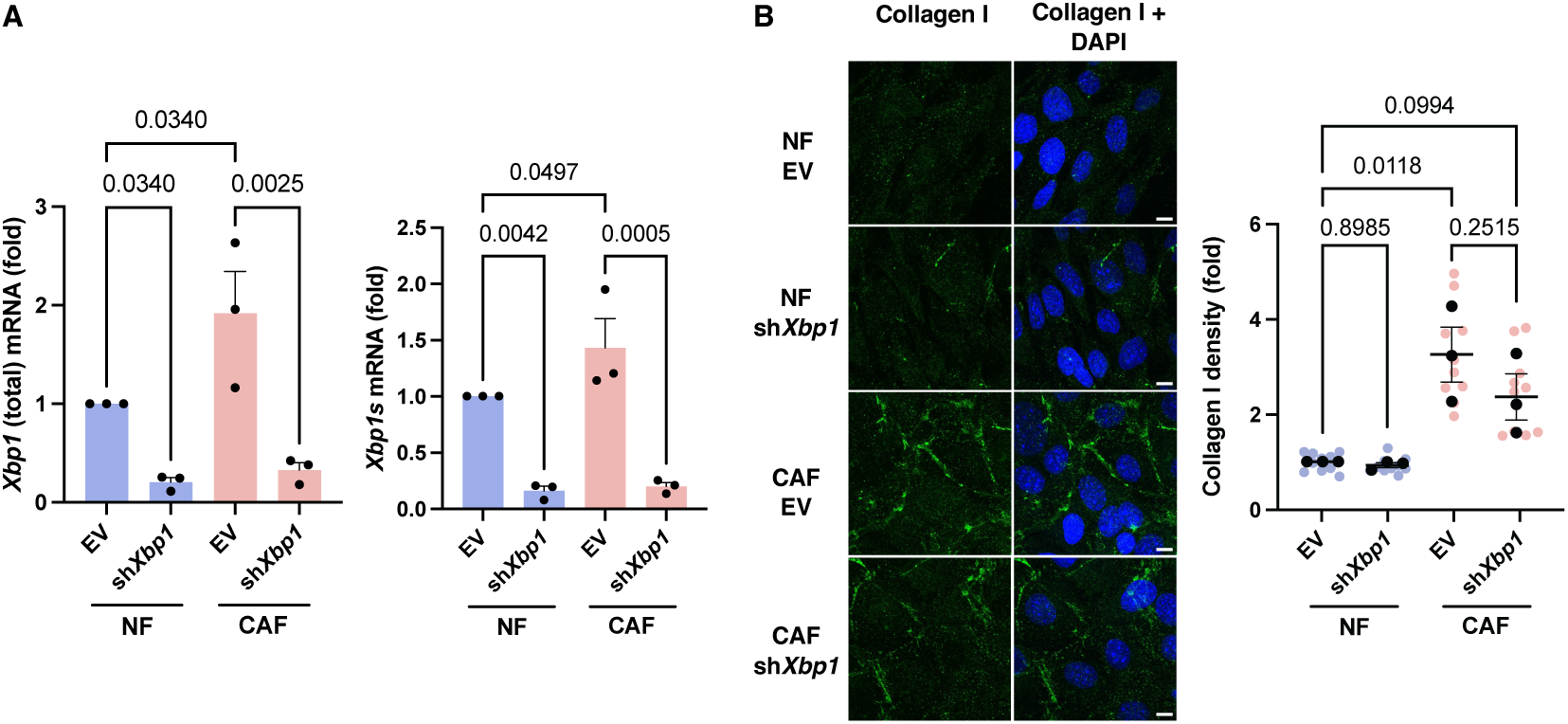
**(A)**  Expression of total *Xbp1* mRNA (left) and spliced *Xbp1* (*Xbp1s*) mRNA (right) in NFs and CAFs transduced with either empty vector (EV) or sh*Xbp1* constructs, n=3. **(B)** IF staining of collagen I (green) in NFs and CAFs transduced with EV or sh*Xbp1* constructs, and quantification of collagen I high-density matrix area normalised to cell number. Nuclei are stained with DAPI (blue). Scale bars represent 10 µm.

**Supplementary Figure 3.**
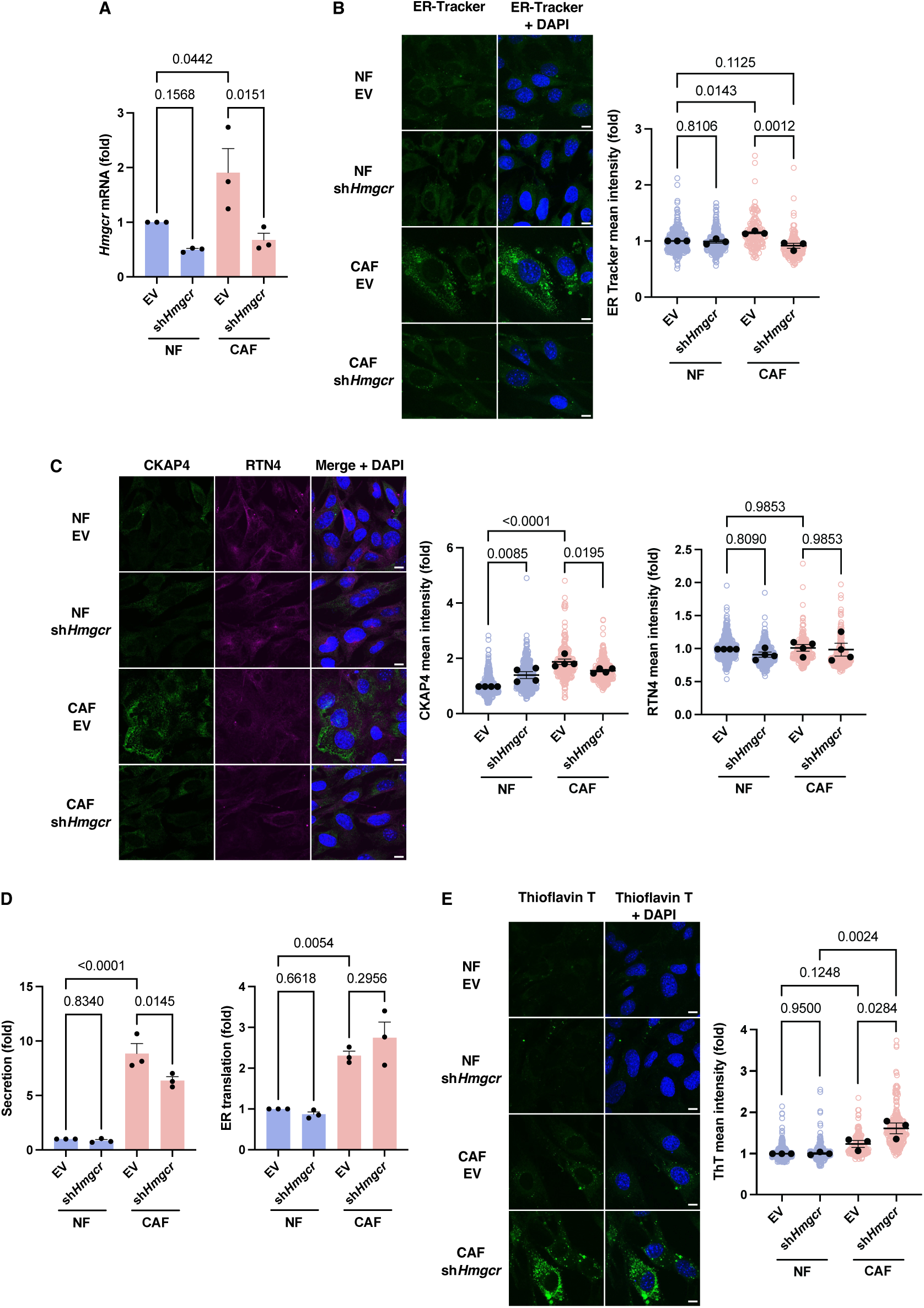
**(A)**  Expression of *Hmgcr* mRNA in NFs and CAFs transduced with empty vector (EV) or sh*Hmgcr* constructs, as determined via qPCR, n=3. **(B)** Representative ER-Tracker staining (green) in NFs and CAFs transduced with EV or sh*Hmgcr* constructs, and quantification of ER-Tracker mean intensity per cell, n=3 independent experiments (NF EV=292 cells, NF sh*Hmgcr*=352 cells, CAF EV=152 cells, CAF sh*Hmgcr*=220 cells). Nuclei are stained with DAPI (blue), scale bars represent 10 µm. **(C)** Representative immunofluorescent staining of ER sheet marker CKAP4 (green) and ER tubule marker RTN4 (magenta) in NFs and CAFs transduced with EV or sh*Hmgcr* constructs, and quantification of CKAP4 and RTN4 mean intensity per cell, n=4 independent experiments (NF EV=482 cells, NF sh*Hmgcr*=474 cells, CAF EV=286 cells, CAF sh*Hmgcr*=352 cells). Nuclei are stained with DAPI (blue). Scale bars represent 10 µm. **(D)** Relative ER translation capacity (intracellular EGFP fluorescence intensity) and relative ER secretory capacity (extracellular GLuc activity) in GLuc-T2A-EGFP-expressing NFs and CAFs transduced with EV or sh*Hmgcr* constructs, n=3. **(F)** Representative ThT staining (green) in NFs and CAFs transduced with EV or sh*Hmgcr* constructs, and quantification of ThT intensity per cell, n=3 independent experiments (NF EV=369 cells, NF sh*Hmgcr* =337 cells, CAF EV=206 cells, CAF sh*Hmgcr* =289 cells). Nuclei are stained with DAPI (blue). Scale bars represent 10 µm.

**Supplementary Figure 4.**
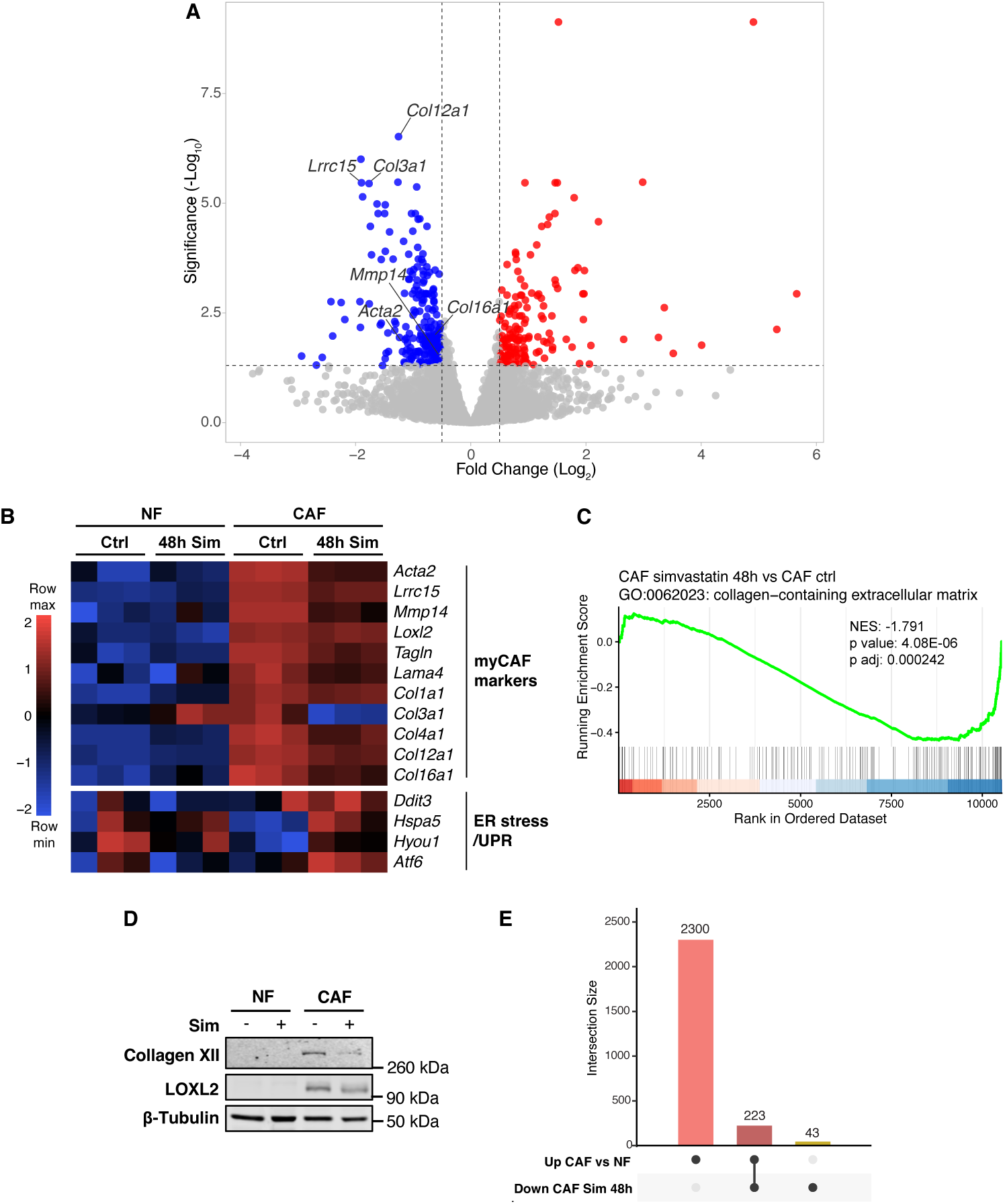
**(A)**  Volcano plot illustrating dinerential gene expression in CAFs following 1 µM simvastatin (Sim) treatment for 48 hours, as determined by RNA-seq analysis. Select genes associated with myCAF function are annotated. **(B)** Heatmap showing relative expression of genes associated with myCAF function and ER stress/UPR in NFs and CAFs treated with DMSO (Ctrl) or 1 µM Sim for 48 hours, as determined by RNA-seq analysis, n=3. **(C)** Representative immunoblot analysis of myCAF-associated marker expression in NFs and CAFs treated with DMSO (Ctrl) or 1 µM Sim for 48 hours. β-tubulin serves as a loading control. **(D)** GSEA plot derived from RNA-Seq analysis of CAFs treated with 1 µM Sim versus CAFs treated with DMSO for 48 hours, showing enrichment of a “collagen-containing extracellular matrix” signature (GO:0062023). **(E)** UpSet plot showing overlap between genes significantly upregulated in CAFs relative to NFs, and those significantly downregulated in CAFs treated with 1 µM Sim for 48 hours relative to CAFs treated with DMSO (Ctrl).

**Supplementary Figure 5.**
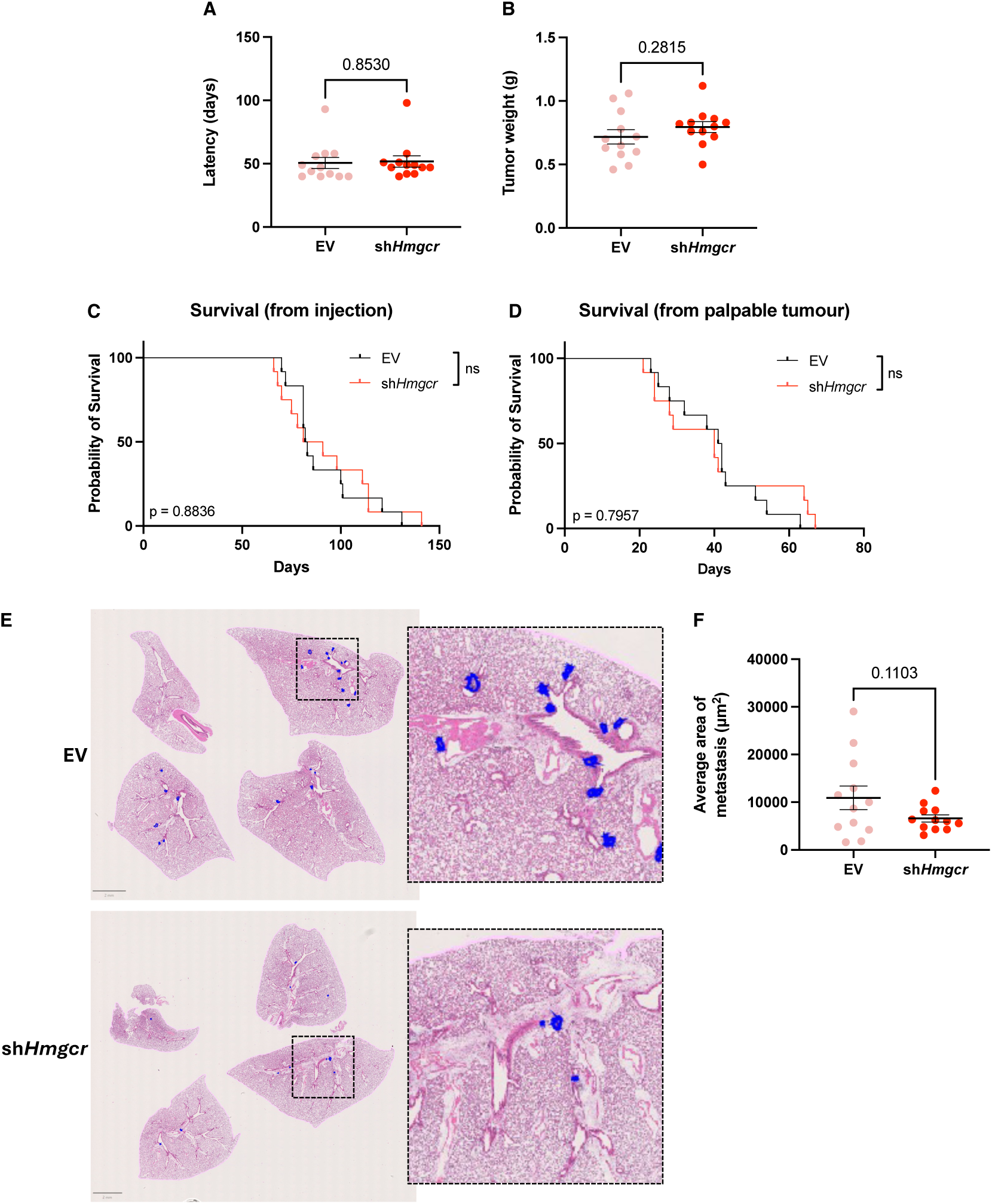
Quantification of **(A)** tumour latency (days from injection to palpable tumour) and **(B)** tumour weight in mice co-injected with PyMT cancer cells and CAFs transduced with EV or *sh*Hmgcr constructs, n=12 mice per group. Kaplan-Meier analysis of **(C)** time from injection to endpoint and **(D)** time from palpable tumour to endpoint of mice co-injected with PyMT cancer cells and CAFs transduced with EV or *sh*Hmgcr constructs, n=12 mice per group. **(E)** Representative images, and zoomed inset images, of H&E-stained lungs from mice co-injected with PyMT cancer cells and CAFs transduced with EV or *sh*Hmgcr constructs. Metastases are outlined in blue. **(F)** Quantification of average lung metastasis area in mice co-injected with PyMT cancer cells and CAFs transduced with EV or sh*Hmgcr* constructs, n=12 mice per group.

## Notes

### Competing Interest Statement

The authors have declared no competing interest.

